# Monomorphic genotypes within a generalist lineage of *Campylobacter jejuni* show signs of global dispersion

**DOI:** 10.1101/054932

**Authors:** Ann-Katrin Llarena, Ji Zhang, Vehkala Minna, Niko Välimäki, Marjaana Hakkinen, Marja-Liisa Hänninen, Mati Roasto, Mihkel Mäesaar, Eduardo Taboada, Dillon Barker, Giuliano Garfolo, Cesare Cammà, Elisabetta Di Giannatale, Jukka Corander, Mirko Rossi

## Abstract

The decreased costs of genome sequencing have increased capability to apply whole-genome sequence on epidemiological surveillance of zoonotic *Campylobacter jejuni*. However, knowledge about how genetically similar epidemiologically linked isolates can be is vital for correct application of this methodology. To address this issue in *C. jejuni* we investigated the spatial and temporal signals in the genomes of a major clonal complex and generalist lineage, ST-45 CC, by exploiting the population structure and genealogy and applying genome-wide association analysis of 340 isolates from across Europe collected over a wide time-range. The occurrence and strength of the geographical signal varied between sublineages and followed the clonal frame when present, while no evidence of a temporal signal was found. Certain sublineages of ST-45 CC formed discrete and genetically isolated clades to which geography and time had left only negligible traces in the genomes. We hypothesize that these ST-45 CC clades form globally expanded monomorphic clones possibly spread across Europe by migratory birds. In addition, we observed an incongruence between the genealogy of the strains and MLST typing, thereby challenging the existing clonal complex definition and use of a common MLST-based nomenclature for the ST-45 CC of *C. jejuni*.

## 1. INTRODUCTION

The use of whole-genome sequencing (WGS) in genomic epidemiology is revolutionizing surveillance and outbreak investigations of bacterial threats to public health. WGS has been successfully used, for example, to limit the spread of nosocomial MRSA (Harris, Cartwright *et al.* 2013), investigate the origin of the Haiti cholera outbreak (Hendriksen, Price *et al.* 2011, Mütreja, Kim *et al.* 2011) and search for signals of host adaption for use in source attribution (Dearlove, Cody *et al.* 2015). WGS is currently used in real-time surveillance of *Listeria monocytogenes* and *Salmonella* Enteritidis by the American Centers for Disease Control and Prevention and the U.S Food and Drug Administration (www.fda.gov) and similar approaches for *E. coli, Campylobacter, Vibrio,* and *Cronobacter,* etc. are expected to come in use in the near future. In addition, both European Food Safety Authority (EFSA) and European Centre for Disease Control and prevention (ECDC) have emphasized the importance of WGS, and advocate the need for transition from classic laboratory methods to WGS in real-time surveillance of infectious diseases (EFSA 2014, ECDC 2015).

*Campylobacter jejuni* is the most common cause of bacterial gastroenteritis worldwide, with an increasing number of cases reported in the European Union (EU) and Finland (Jaakola, Lyytikäinen *et al.* 2015, EFSA and ECDC 2015). As most cases are self-limiting and unreported and since large point-source outbreaks are rare, identifying sources of *C. jejuni* is difficult (Blaser, Engberg 2008). As a result, 30 years of intense research on *C. jejuni* and various mitigation strategies have not been able to reduce the health burden of campylobacteriosis. Improved methods to attribute sporadic cases and detect hidden outbreaks are needed, and thus considerable expectations are directed towards WGS in this regard to ultimately prevent and control the *Campylobacter* epidemic. Applications of WGS for public health purposes are dependent on knowledge of the genomic relationships between isolates, both in the context of outbreaks and sporadic cases. Also, knowledge on potential genomic changes occurring through a transmission pathway such as the food-chain will be essential in source attribution. According to previous studies on the genetic relatedness of *C. jejuni* circulating in outbreaks and clustering in time and space in chickens, genetic diversity varies between multilocus sequence types (ST) and clonal complexes (CC) (Revez, Llarena *et al.* 2014, Revez, Zhang *et al.* 2014, Kivistö, Kovanen *et al.* 2014, Kovanen, Kivistö *et al.* 2016). Such differences between lineages and sublineages complicate the development of an universal nomenclature, and more studies on the genetic diversity within and across lineages is warranted.

ST-45 CC is a generalist lineage having a wide range of host animals (Sheppard, Cheng *et al.* 2014) and a rapid host-switch rate that appears to erode signs of host-adaption (Dearlove, Cody et al. 2015). Within this CC, the founder ST-45 is very heterogeneous by Penner heat-stable serotyping (Dingle, Colles *et al.* 2001), *flaA* short-variable-region typing (Dingle, Colles *et al.* 2002), comparative genomic hybridization (Taboada, Mackinnon *et al.* 2008), stress response analyses (Habib, Louwen *et al.* 2009), lipooligosaccharide locus class distributions (Revez and Hänninen 2012), and whole genome multilocus sequencing typing (Kovanen, Kivisto *et al.* 2016). The existence of several possible animal host species, lack of host signals and genetic and phenotypic heterogeneity of this CC complicate the use of WGS in epidemiological investigations and source attribution. Therefore, a robust description of the genomic diversity within the ST-45 CC across time and space, the two main factors relevant in public health surveillance, is essential to better understand the genetic relationship between two isolates.

Genetic differences within species due to geography are commonly encountered in prokaryotes, for example *Helicobacter pylori* (Linz, Balloux *et al.* 2007). In *C. jejuni,* this phenomenon is reflected in the overrepresentation or exclusiveness of different lineages according to geography, such as ST-474 in New Zealand (Müllner, Collins-Emerson *et al.* 2010) and ST-677 in Finland (de Haan, Kivisto *et al.* 2010, Kärenlampi, Rautelin *et al.* 2007). Although members of ST-45 CC have been isolated worldwide, there is a high relative frequency of this CC among Finnish patients and chickens compared to other countries (de Haan, Kivisto *et al.* 2010, McCarthy, Gillespie *et al.* 2012, Llarena, Huneau *et al.* 2015). However, due to the limited resolution of MLST, it is unclear whether the Finnish overrepresentation of this lineage is a consequence of the local expansion of a successful clone with limited geographical distribution. In this concern, WGS and genome wide association studies (GWAS) have the potential to provide insight on the process of evolution that have favoured one lineage over another.

The degree of genetic diversity over time varies considerably between prokaryotes, which has strong implications for the applicability of WGS in pathogen surveillance. For instance, *Yersinia pestis* has been under strong purifying selection and has been nearly unaltered since the Black plague (Achtman 2012), while, in opposite, Morelli *et al.* (2010) found at least 124 single nucleotide variants over a 40 000 nt region accumulated during a decade in *H. pylori.* Neither long term evolutionary studies nor studies on the evolutionary change over time in a natural population setting are currently available for *C. jejuni.* However, Wilson *et al.* (2009) proposed an absolute mutation rate for *C. jejuni,* calculated from MLST, of 3.23 × 10^−5^ substitutions per site per year. This estimation is ten times faster than the one calculated for *H. pylori* (Morelli, Didelot *et al.* 2010) and *Pseudomonas aeruginosa* during chronic infections (Smith, Buckley *et al.* 2006), and hundred times faster than estimates for *E. coli* (Reeves, Liu *et al.* 2011). Therefore, without ignoring the limitations of these estimates (Morelli, Didelot *et al.* 2010, Kuo, Ochman 2009), assuming Wilson’s clock rate and, hence, consequently the predicted time of divergence of the most recent common ancestor of ST-45 CC (~81 years before present; Dearlove *et al.,* 2015), detectable evolution and separation by time is expected over the due course of a decade.

Our main aim was to characterize the variation and diversity in ST-45 CC across time and space. By comparing 340 isolates of British, Finnish and Baltic origin, we searched for spatial and temporal signals in the genomes of ST-45 CC isolates with the ultimate aim to evaluate the applicability of WGS analysis in surveillance and outbreak investigations. We sought to answer the following two questions: how heterogeneous are various ST-45 CC sublineages and how, if at all, do the genomes of this clonal complex vary over time and between countries?

## MATERIALS AND METHODS

### Isolates, genome sequencing and assembly

The genomes of 199 *C. jejuni* ST-45 CC isolates collected between 2000 and 2012 in the United Kingdom (UK), including isolates collected through the Oxfordshire sentinel surveillance study (http://pubmlst.org/campylobacter/info/Oxfordshiresentinelsurveillance.shtml), were selected from the PubMLST database (http://www.pubmlst.org/; accessed May 2015 (Jolley, Maiden 2010)). Furthermore, the sequenced genomes of 125 *C. jejuni* ST-45 CC isolates of Finnish origin from 2000 to 2012 were included from earlier studies (Revez, Llarena *et al.* 2014, Kovanen, Kivisto *et al.* 2014b, Llarena, Huneau *et al.* 2015, Zhang, Halkilahti *et al.* 2015). Also, 17 *C. jejuni* ST-45 CC isolates of Finnish, Estonian and Lithuanian origin from 1999 to 2012 (Kärenlampi, Rautelin *et al.* 2007, Kovanen, Kivistö *et al.* 2014a) and one *C. jejuni* ST-21 CC isolate from Estonia 2012 were whole-genome sequenced using Illumina technology (performed by FIMM, University of Helsinki, Finland). Genome assembly was performed as described earlier (Kovanen, Kivistö *et al.* 2016). This dataset included a total of 22 STs from ST-45 CC, of which 13 were considered singleton STs as they accounted for two or fewer isolates. All genomes included in this study were smaller than 1.8 Mb and assembled with ≤ 100 contigs and the entire isolate collection and metadata are presented in Table 1 and S1 with necessary PubMLST database accession numbers.

**Table 1:** Overview of isolates, their hosts and country of origin included in the study.

| Source | Number | Country | Source of genome sequence |
| --- | --- | --- | --- |
| ST-45 CC |  |  |  |
| Human | 144 | UK | PubMLST |
|  | 36 | Finland | Kovanen, Kivistö et al. (2014) |
| Animal | 42 | UK | PubMLST |
|  | 95 | Finland | Llarena, Huneau et al. (2015),<br>Zhang, Halkilahti et al. (2015),<br>this study |
|  | 3 | Estonia | This study |
|  | 2 | Lithuania | This study |
| Environment (water) | 5 | Finland | Kovanen, Kivistö et al. (2016) |
| Unknown | 13 | UK | PubMLST |
| ST-21 CC |  |  |  |
| Animal | 1 | Estonia | This study |

### Generation of the core and accessory genome

Prodigal was used for gene prediction and a database including all translated coding sequences (tCDSs, n = 581,171) of the *C. jejuni* genomes was assembled. Reciprocal all-versus-all BLASTp search was performed (threshold *E ≤*1e^−10^) (Altschul, Madden *et al.* 1997) and orthologous groups were determined by orthAgogue and MCL (ignoring E-values, percent match length ≥ 80% and inflation value of 1.5) (Enright, Van Dongen *et al.* 2002, Ekseth, Kuiper *et al.* 2014). The groups of orthologs (GOs) were aligned using MUSCLE and back-translated to nucleotide sequence using Translatorx perl script (Edgar 2004a, Edgar 2004b, Abascal, Zardoya *et al.* 2010). GOs with a total alignment length less than 300 bp. were excluded. Core and accessory gene pools were extracted and one representative sequence from each of the GOs from the accessory genome was stochastically selected and annotated using Rapid Annotation Server (Aziz, Bartels *et al.* 2008). This annotation was transferred to all members of the corresponding GO. The presence of plasmids (pVir and pTet) and integrated elements (CJIE1-5) (Batchelor, Pearson *et al.* 2004, Fouts, Mongodin *et al.* 2005, Hofreuter, Tsai *et al.* 2006, Skarp, Akinrinade *et al.* 2015) were inferred using a customized perl script (available on request). Genes and genetic structures suggested to be involved in niche adaption or strain variability (Hofreuter, Novik *et al.* 2008, Sheppard, Didelot *et al.* 2013, Vorwerk, Huber *et al.* 2015), including metabolism and antibiotic resistance loci, were selected for further analysis (e.g. association with population structure), while hypothetical and weakly annotated proteins were excluded.

### Population genetics, genealogy reconstruction and pangenome analysis

Population structure was defined using BAPS 6.0 (the module hierarchical BAPS (hierBAPS)) with default settings (Corander, Waldmann *et al.* 2003, Cheng, Connor *et al.* 2013). The analysis was performed on a fraction of the concatenated multisequence alignment of *C. jejuni* ST-45 CC strains with orthologs in the *C. jejuni* ST-21 strain (1043 GOs). The number of base differences within BAPS clusters was calculated in MEGA5 (Tamura, Peterson *et al.* 2011) based on pairwise base differences averaged over all possible sequence pairs.

For genealogy reconstruction, recombination was identified and excluded from the concatenated core-genome defined above using BratNextGen with 20 iterations of the HMM estimation algorithm and 100 permutations with 5% significance threshold (Marttinen, Hanage *et al.* 2012). Thereafter, phylogeny was inferred using a maximum likelihood (ML) tree based on the recombination-free core-genome sequence estimated with RAxML v. 7 under the generalized time-reversible model (GTRGAMMA) with 100 bootstrap replicates (Stamatakis 2014). Moreover, a pangenome binary matrix was built for all strains and used as input to an ML tree construction by RAxML, applying the evolutionary model for binary data with 100 bootstrap replicates. Both ML trees were rooted at the split of the outgroup strain. In addition, the isolates metadata, population structure and the presence or absence of selected genes were colour-coded to both ML trees using iTOL v 3.0. (Ciccarelli, Doerks *et al.* 2006).

## Validation

The genomes of five ST-45 strains isolated from barnacle geese in a previous study (Llarena *et al.,* 2015) were sequenced as described earlier (Kovanen, Kivisto *et al.* 2016). To assess the global distribution of two clonal sublineages of ST-45 and whether migrating birds could participate to the dissemination of these across country borders, the genomes of 101 and 8 ST-45 isolates of Canadian and Italian origin, respectively, were compared with one representative of each hierBAPS cluster level 2 on lineage *b* and the five barnacle geese isolates using Parsnip software (Treangen, Ondov *et al.* 2014).

## GWAS and phylogeographic analyses

To identify genetic signals overrepresented in strains originating from Finland and the Baltic countries (analysed as a group designated “Baltic”) or the UK, GWAS was performed using sequence element enrichment analysis as described in Weinert *et al.* (2015), with the minimum and maximum length of k-mers equal to 10 and 100, respectively. The population structure was accounted for by the hierBAPS clusters and the significance threshold used for any single k-mer was 10^-8^. An intensity plot was created by mapping the significant k-mer frequency per 100 bp. along the reference genomes of *C. jejuni* M1 (Friis, Wassenaar *et al.,* 2010) and 4031 (Revez, Llarena *et al.* 2014) for the isolates of British and Baltic origin, respectively. A cut-off value for significant k-mer frequency of five was used to identify regions of interest. To analyse the distribution of these words in British and Baltic isolates according to population structure, open reading frames of each region were extracted from the reference genome and used to assess the presence of the three regions A, B and C in the dataset using a gene-by-gene approach implemented in Genome Profiler (GeP) (flag-o, Zhang *et al.,* 2015). Missing or truncated loci were considered non-present. The association between these regions and the isolates' geographical origin was assessed using Pearson's Chi-Square and Fisher exact test within level one BAPS clusters *(P* < 0.05 considered significant).

## Timescale of evolution of *C. Jejuni*

To estimate the genetic variance across time, Finnish isolates collected during 13 years grouping to BAPS clusters 2, 4, and 6 were selected. For each BAPS cluster, the non-recombined core-genome alignment was extracted using GeP v 2.0 (Zhang, Halkilahti *et al.* 2015) and BratNextGen (Marttinen, Hanage *et al.* 2012). Finally, the alignment gaps were removed, resulting in a gap-free core genome alignment of approximately 1.2 – 1.3 Mb. A dated phylogeny of each BAPS cluster was reconstructed using Bayesian Evolutionary Analysis Sampling Trees (BEAST) v.1.8.0 (Drummond, Suchard *et al.* 2012) assuming a constant population size and with prior parameters normally distributed (mean 0.0; SD 1.0). The HKY substitution model with a strict evolutionary clock was used (Hasegawa, Kishino *et al.* 1985, Drummond, Bouckaert 2015), implying a uniform rate of evolution across branches. For the evolutionary clock, a log normal prior (mean −10.0, SD 1.0, initial value 3.0 × 10^−5^) was used. The Markov Chain Monte Carlo was run with 5 million iterations, 500 000 burn in iterations, and sampling every subsequent 1000 iterations. The chain was twice and summarized in one BEAST tree for each BAPS cluster. An exploratory root-to-tip linear regression, using sampling year information for each strain as independent variable, was performed in TempEst v1.4 on each Bayesian tree (Rambaut, Lam *et al.* 2016). Thereafter, the coalescent time (denoted τ) estimated by BEAST was tentatively calibrated to years as described earlier (Dearlove, Cody *et al.* 2015) using the Wilson absolute mutation rate for *Campylobacter* of 3.23 × 10^−5^ substitutions per site per year (Wilson, Gabriel *et al.* 2009).

## Statistics

The isolates spatial association with population structure was tested by Pearson Chi-Square or Fisher exact-test. Furthermore, correlation between number of STs, years, and countries of origin and genetic distance was calculated. All statistics were done in SPSS v. 5.1 (International Business Machines Corp., Armonk, NY, USA) with *P* < 0.05 considered significant.

## RESULTS

### Population structure and core genealogy

The core genome was defined as the GOs present in at least 99% of the 340 isolates, and contained 1383 of the total 2664 GOs. GOs with orthologs in the *C. jejuni* ST-21 strain (n = 1043) were used to reconstruct genealogy and infer population structure (phylogram Fig. 1A; inner phylogram Fig 1B; cladogram with bootstrap supporting values ≥ 80% in Fig S1A). Two major clades were identified *(a* and *b,* Fig. 1 and Fig. S1), and the most frequent STs in the dataset (ST-45, ST-230, ST-137, and ST-583) grouped together on clade b. Recombination was more extensive in clade *a;* on average 70.1% (r/m ratio ~22) and 8.4 % (r/m ratio ~3) of the core genome was recombining in *a* and b, respectively.

**Figure 1A.**
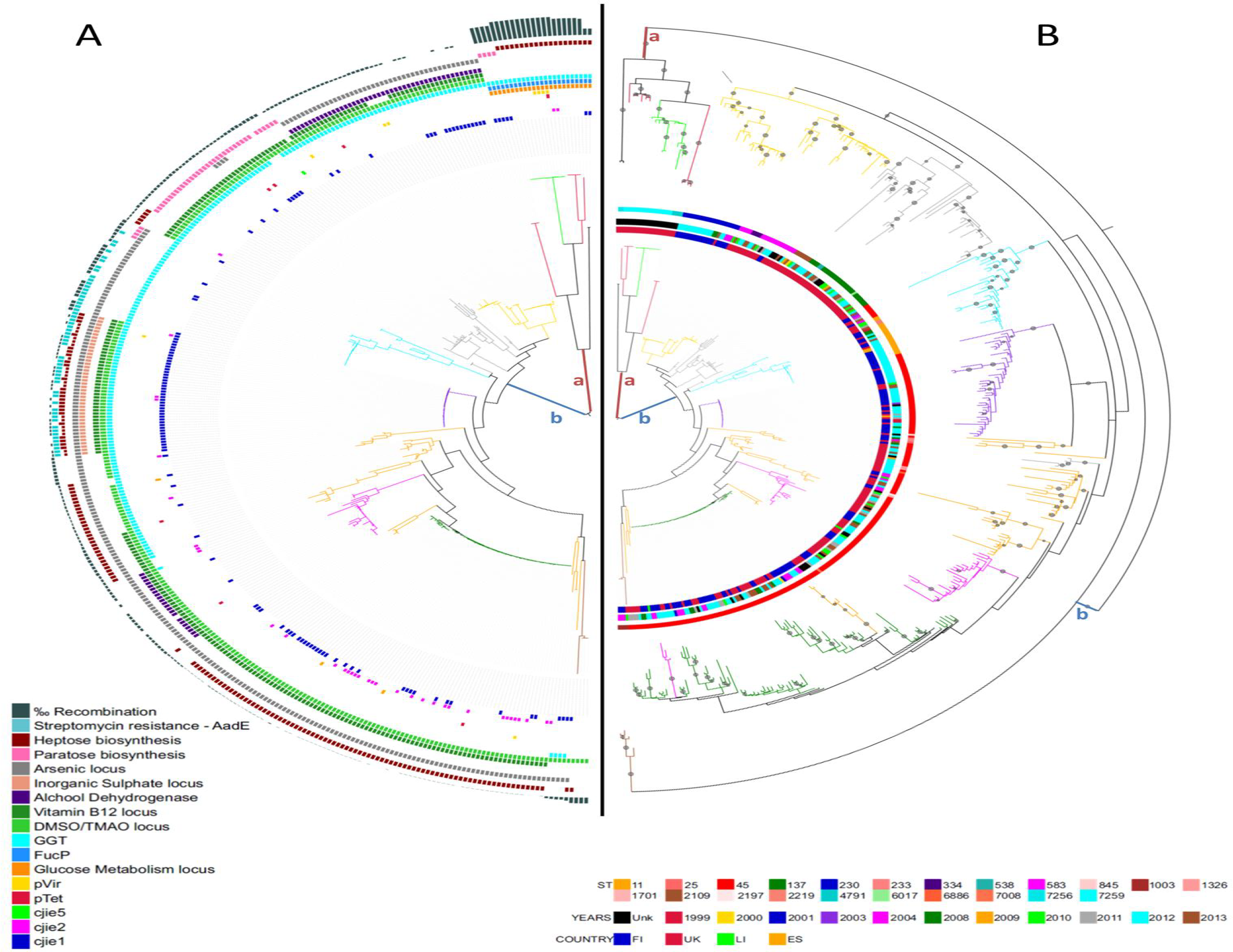
Phylogram of RAxML tree based on non-recombined core genome. Semicircles from out to in represented genetic features listed in the legends. Branches are colored according to hierBAPS clustering. The two major clades are indicated with letters *a* and *b.* Figure 1B: *Inner phylogram:* RAxML tree based on unrecombined core genome as described in 1A. *Outer phylogram:* RAxML tree based on binary matrix of presence-absence of accessory genes. Nodes with bootstrap values > 50% are indicated with a circle. Semicircles from out to in: STs, years, origin, and indications of which are in the legends. Branches are colored according to hierBAPS clustering. See text for further details.

Population structure was inferred using nested clustering implemented in hierBAPS and details are presented in Table 2. First level hierBAPS divided ST-45 CC between 11 BAPS populations (BAPS 1, 2, 4–12) which split further into 38 subpopulations on the secondary BAPS level (BAPS 1^*^–7^*^, 9^*^–38^*^, star denounces level 2). BAPS 1, 8 and 12 were on clade *a*, while the remaining BAPS groups were polychotomously arranged (bootstrap values ≥ 80%) in six monophyletic groups within clade *b* (Fig 1A, Fig S1A, Table S2). Except for the polyphyletic ST-45 and paraphyletic ST-7259, first level hierBAPS did not divide STs into different populations (Table 2 and S2). At secondary level, 243 isolates belonged to poly- or paraphyletic STs which split into different BAPS^*^ clusters, e.g. ST-45 and ST-7259 split in 18 and 5 subpopulations on the second BAPS level, respectively (Table 2 and S2). Four STs (ST- 230, ST-334, ST-2109 and ST-4791, 37 isolates) were monophyletic and formed ST-specific populations.

**Table 2.** Overview of population structure of ST-45 CC with associated geographical signals according to clonal frame. Geographic association is illustrated in bold letter with respective country or region.

| BAPS L1 | bp. diff.<br>* | n** | STs | Countries | Year | Origin | Phyly# | Recom#<br># |
| --- | --- | --- | --- | --- | --- | --- | --- | --- |
| Clade *a*& |  |  |  |  |  |  |  |  |
| 1& | $1,7 \cdot 10^{-2}$ | 10 | ST-4791, ST-7259 | UK | UNK | A | Para | 0,701 |
| 8& | $7,4 \cdot 10^{-5}$ | 12 | ST- 7259 | UK | UNK | A | Mono | 0,804 |
| 12& | $2,7 \cdot 10^{-6}$ | 2 | ST- 7256 | UK | UNK | A | Mono | 0,262 |
| Clade *b* |  |  |  |  |  |  |  |  |
| 2& | $1,5 \cdot 10^{-3}$ | 46 | ST-230 (FIN), ST-334, ST-583 (UK) | FIN, UK | 2004-2013 | A, H, E | Mono | 0,059 |
| 4 | $2,4 \cdot 10^{-5}$ | 42 | ST- 45 | EST, FIN, UK | 1999-2013 | A, H, E | Mono | 0,042 |
| 5 | $1,0 \cdot 10^{-5}$ | 26 | ST- 45 | LIT, FIN, UK | 2008-2013 | A, H, E | Mono | 0,075 |
| 6 | $9,2 \cdot 10^{-5}$ | 83 | ST- 45, ST-7008 | LIT, FIN, UK | 2000-2013 | A, H, E | Mono | 0,001 |
| 7& | $4,4 \cdot 10^{-3}$ | 45 | ST-25, ST- 45, ST-233, ST-686, ST-845, ST-1326, ST-1701, ST-2197 | FIN, UK&& | 2004-2013 | A, H | Poly | 0,054 |
| 9& | $3,4 \cdot 10^{-3}$ | 42 | ST-45, ST- 137, ST-538, ST-2109, ST-6017 | FIN, UK | 2000-2013 | A, H | Poly | 0,078 |
| 10& | $1,6 \cdot 10^{-3}$ | 28 | ST- 11 (BAL), ST-45, ST-2219 (BAL) | BAL, UK | 1999-2013 | A, H | Mono | 0,124 |
| 11 | $5,1 \cdot 10^{-3}$ | 4 | ST- 1003 | UK, FIN | 2004-2010 | A, H | Mono | 0,237 |

\* Average genetic distance nucleotide in the aligned, non-recombining core genome
\*\*number of strains in each BAPS cluster

The average base differences per nucleotide in the non-recombined core genome across population structure ranged between 1.7–10^−2^ and 2.7–10^−6^ (Table 2). Higher genetic diversity (1.7–10^−2^ to 1.5–10^−3^) was observed in BAPS clusters populated by multiple STs (BAPS 1, 2, 7, 9 and 10) reflected in a positive correlation between genetic distance and diversity of STs in the BAPS groups (0.693; p=0.018, and 0,722; p=0,012). Low genetic difference was evident for major BAPS groups, especially BAPS 4 (Fig. 1A; purple clade), 5 and 6 (Fig. 1A; green clade) composed of ST-45.

The geographical signal varied between branches in the core genealogy and BAPS clusters (Table 2). Clade *a* was British, while BAPS 2 on clade *b* divided between one Finnish and one British branch with ST-230 and ST-583/ ST-334, respectively. BAPS 9 was British, except for two Finnish isolates in sub-population BAPS 32^*^. The geographical structure of the polyphyletic BAPS 7 population varied with BAPS level 2; 22^*^– 24^*^ were mostly British while 20^*^ and 21^*^ were of mixed origin. In BAPS 10, branches containing strains of ST-11 and ST- 2219 were all of Baltic and Finnish origin, while in the remaining ST-45 strains from the UK and the Baltics were equally distributed between the branches. BAPS 4 was overrepresented by Finnish strains, but 25% of the isolates were British and the clonal genealogy was unable to separate the strains due to low genetic distance. Similarly, strains of BAPS 5, 6, and 11 did not group according to geography. Most of the strains (95.8%) in BAPS populations lacking geographical signal was ST-45.

Isolates from a wide range of sampling times were scattered through the ML tree (Fig. 1), with no evidence of monophyletic relationship in terms of time. The population structure could not predict the temporal origin of the isolates in a clear-cut way as all BAPS populations on level 1 and 2 contained isolates from different years (for BAPS populations above five isolates; Table 2). This lack of genetic structuring in terms of time was reflected in the genetic distance which was not associated with the number of sampling years covered by the BAPS populations.

### Pangenomic analysis

To assess the relationship between the accessory- and core genomes and the population structure and to search for potential spatial and temporal signals residing in the accessory genome, a ML tree based on the presence and absence of genes was built. The branch lengths increased considerably with the inclusion of the accessory genome and a clearer separation of isolates inside BAPS 4, 6, and partially BAPS 10 was seen (outer phylogram Fig. 1B). The increased resolution offered by this analysis revealed low levels of geographical clustering inside BAPS 6 which was not apparent in the core genome analysis, as a few Finnish and British clades emerged, possibly due to geographically local microevolution. No further geographical structuring was evident in the remaining clusters or populations. Again, no temporal signals were found in the accessory genome as no grouping according to sampling year was seen. The inclusion of the accessory genome decreased the support for the deepest branches of the tree substantially, probably due to the occurrence of pervasive horizontal gene transfer (bootstrap values ≥ 80%, Fig S1B). The two major clades, *a* and b, were still present and the constituents of these two lines remained. However, the polychotomy of clade *b* increased drastically, splitting the six core genealogy-based monophyletic groups into 73, i.e. BAPS 6 was divided into 57 single lineages. Increased polychotomy was also evident for BAPS 4 and 7. Moreover, BAPS 5 changed from a mono- to a paraphyletic lineage, and BAPS 2 and BAPS 9 split and thereby making BAPS 9 nearly monophyletic for UK origin.

An association between specific sets of accessory genes and the clonal frame was found across the CC (Fig 1A). Clade *a* contained strains characterized by the unique combination of genes for glucose and fucose metabolism *fucP)* and the gamma-glutamyl transferase gene *(ggt).* On clade b, each monophyletic group was characterized by a specific assortment of genes related to metabolism (Vitamin B5 biosynthesis, *ggt,* alcohol dehydrogenase, arsenic, inorganic sulphate and DMSO/TMAO metabolism), surface structures (heptose and paratose biosynthesis), hypervariable regions *(AadE,* CJIE1-5) and plasmids (pTet and pVir).

### Identification of phylogeographical signal in ST-45 CC isolates using GWAS

By the use of GWAS, three regions in the accessory genome (marked A, B and C) were identified as overrepresented in strains originating from either the UK or Finland, Estonia and Lithuania, of which the three latter were analysed as a group. The Manhattan plots are presented in Fig. 2, and show that region A and B were enriched in British strains, while Region C was overrepresented in the Finnish and Baltic strains. Regions A and B matches the C-terminal parts (~100 nt) of two identical methyl-accepting chemotaxis protein paralogs and Region C corresponds to a hypothetical protein. As the impact of these signals was uncertain, the distribution of these signals in British and Finnish-Baltic strains according to population structure was assessed. Except for BAPS 4, 5, and 6, in which Region A and B were associated with British isolates, none of the three signals were able to correctly predict geographical origin within populations.

**Figure 2.**
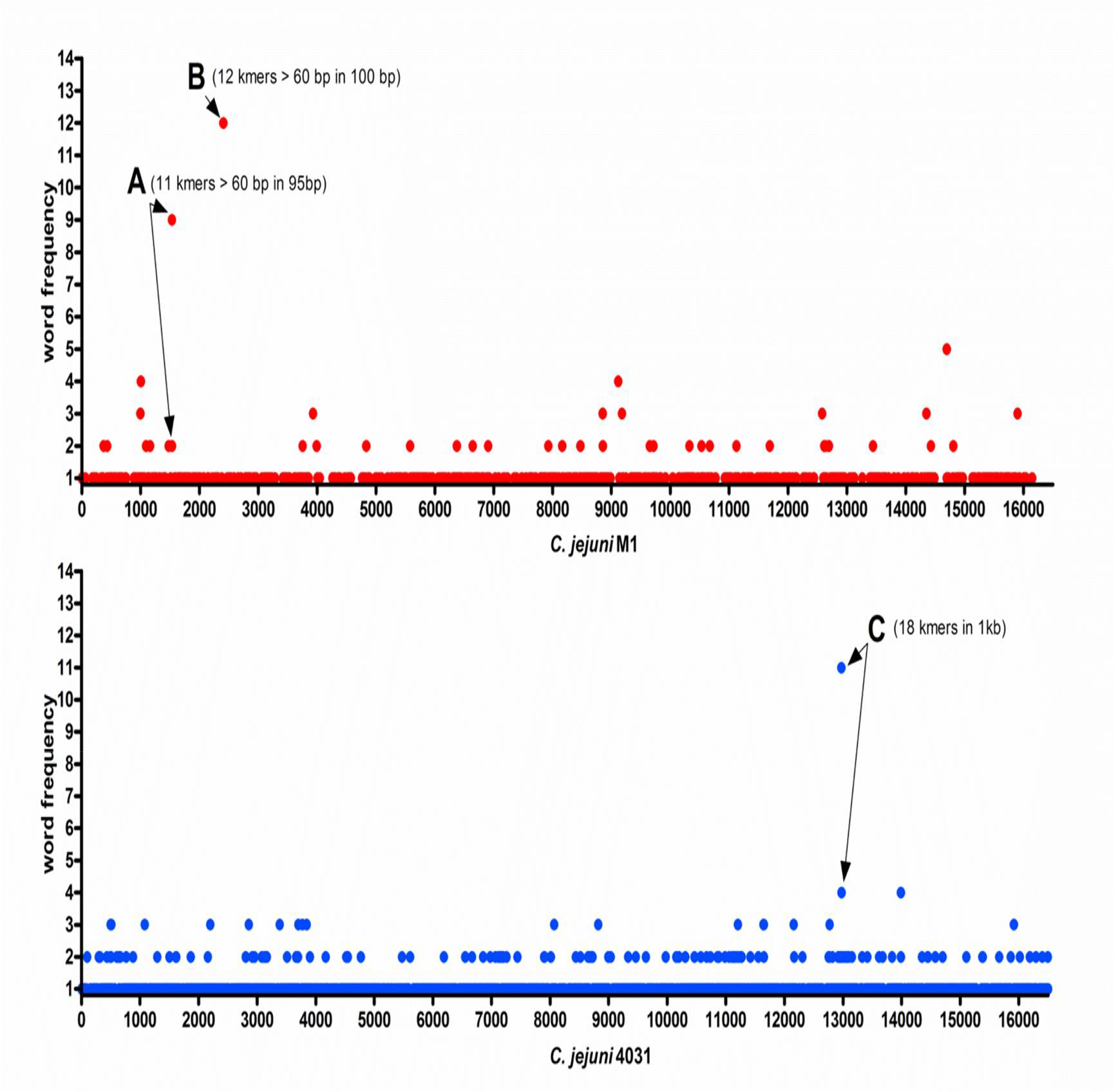
Manhattan plots showing the significant hits associated to either UK (red, upper) or Finland/Baltic countries (blue, lower) mapped to *C.jejuni* M1 or *C. jejuni* 4031 genomes, respectively. The dots represent number of words (kmers), and the regions with higher number of mapped words are indicated with arrows and letters. In the upper UK Manhattan plot, region A of 11 kmers mapped to position 153,200 – 153,295 while region B of 12 kmers at position 240,500 – 240,600, both on strain M1. In the lower Finnish/Baltic plot, region C of 18kmers mapped to position 1,276,300 – 1,277,300 on the 4031 reference genome. See text for further details.

### Evidence for monomorphic clones in ST-45 CC

The geographical signals in the accessory- and core-genome of ST-45 CC varied between sublineages, implying variation in the occurrence of local microevolution. Such lack of genetic isolation by distance might be due to the monomorphic nature of certain lineages which are more genetically stable over time and space (Achtman 2012). We therefore investigated the occurrence of a temporal signal using BEAST for Finnish isolates within three populations harbouring low genetic diversity in their core genome; two ST-45 populations (BAPS 4 and 6) and one ST-230 sub-lineage (BAPS 2). Three separate clades were identified in BAPS 6 (Fig 3), and one of them was populated by strains collected during the whole sampling period. In this clade, the nucleotide diversity in the non-recombined core-genome ranged between 0 and 83, which was similar to the average nucleotide diversity measured between isolates collected during one year (as highlighted in Fig 3). Using a molecular clock with the mutation rate 3.2 × 10^−5^ nucleotides per year (Wilson, Gabriel *et al.* 2009), the time of the most common recent ancestor for this cluster was estimated to be 1.1 years before present (node B in Fig. 3A), which corresponds poorly with the true sampling points of these strains. In addition, the sampling date had a weak correlation with the root-to-tip distance calculated in TempEst v1.4 (linear regression, R^2^ = 0.12). Similar results were observed in ST- 230 and BAPS 4 (data not shown), suggesting that these *C. jejuni* ST-45 CC populations are genetically monomorphic sub-lineages evolving at a much slower rate than previously estimated.

**Figure 3.**
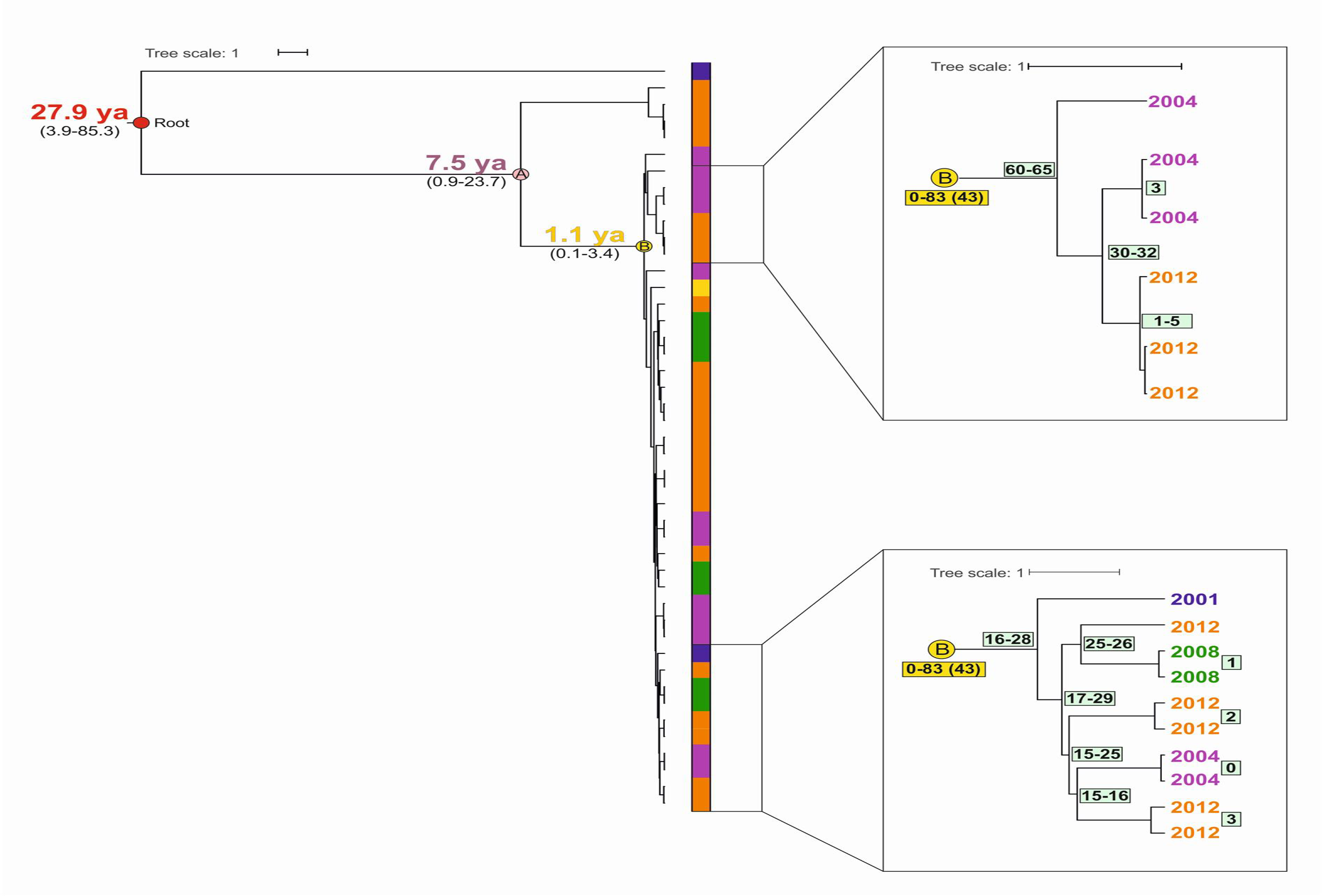
Bayesian phylogenetic tree of Finnish BAPS 6 isolates inferred with BEAST based on unrecombined core genome of ~1.2 Mb. Year of isolation is indicated in the color-bar: blue: 2001, purple: 2004, green: 2008, orange: 2012, yellow:, dark blue: Two subclusters of the clonal BAPS 6 population are scaled up and presented as two embedded illustrations. Predicted divergence times with confidence intervals given in years before present are given at their respective nodes. Single nucleotide variant numbers in a cluster are given in yellow and light green boxes by the most common recent ancestor of that cluster (when space allows, otherwise the box is located between the involved leafs). Fewer than 5 single nucleotides variant were detected between isolates epidemiologically linked (two from 2008, two from 2012, two from 2004, and three from 2012), confirming the usefulness of WGS in outbreak investigations. See main text for further details.

### The disseminator role of migratory birds and global distribution of monomorphic ST-45

To assess if migrating birds could participate in the dissemination of *C. jejuni* across borders, the genomes of *C. jejuni* ST-45 isolates obtained from barnacle geese were sequenced and mapped on the ST-45 CC core tree. One and two geese isolates clustered within the clonal clades of BAPS 6 and 4, respectively (Fig. 4), indicating possible transport of these monomorphic genotypes by migratory birds between Finland and UK.

**Figure 4.**
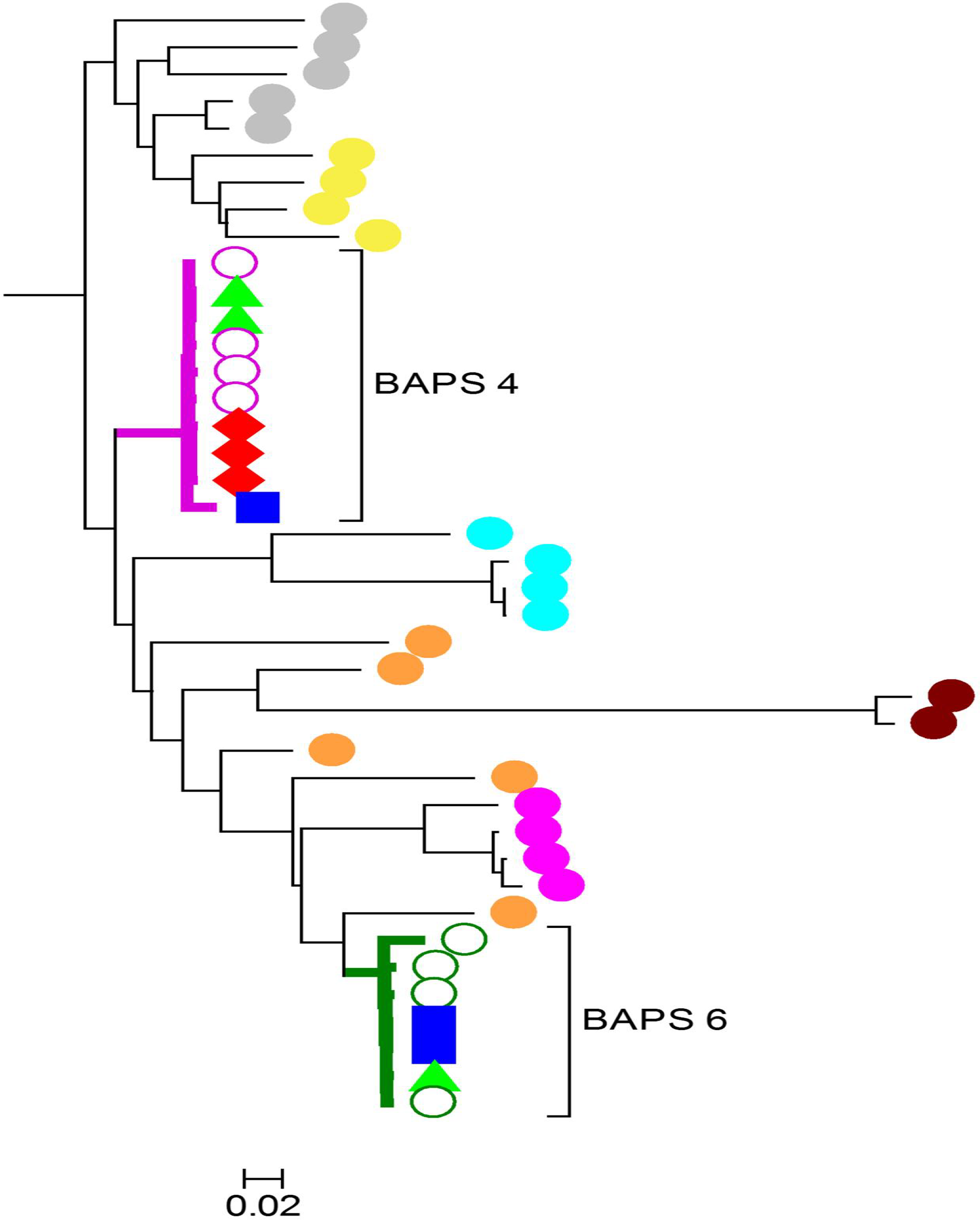
Comparison of 106 ST-45 of Canadian, Italian and Barnacle geese origin and one representative of each hierBAPS cluster level 2 on the *b*-lineage. Purple open circles: Baps 4, green open circles: BAPS 6, Green triangle: barnacle geese, red combo: Canadian strain, blue squares: Italian. BAP 4 and 6 are highlighted for clarity. Genetic distance is indicated with a bar. See main text for further details

To assess if the dissemination of monomorphic clones was limited to the sampled geographical regions, the genomes of *C. jejuni* ST-45 CC isolates of Canadian and Italian origin were mapped against ST-45 CC core genome. Three Canadian and one Italian *C. jejuni* isolated from chickens in 2011 and 2015, respectively, clustered within the diversity of the clonal BAPS 4 cluster. Moreover, two Italian chicken *C. jejuni* isolates from 2015 clustered with the BAPS 6 population (Fig. 4). Overall, these data suggest a global dispersion of these monomorphic *C. jejuni* genotypes.

## DISCUSSION

Our study on the genomic diversity in a *C. jejuni* lineage across space and time is novel and of crucial important in the assessment of molecular epidemiology of campylobacteriosis in the context of public health. By using various state-of-the art methods to search for spatial and temporal signals in a representative strain collection of the widespread ST-45 CC, we found an overrepresentation of certain sub-lineages in the UK and Finland. However, the presence of the geographical signals varied between these and, when present, followed the clonal frame. Contrary to this, geography and time had no effect on genetic diversity of isolates in two BAPS populations of ST-45. Therefore, we propose that these two ST-45 sublineages are globally circulating monomorphic clones, possibly disseminated by migratory birds as suggested by the additional data of strains from barnacle geese. Secondly, we present evidences that challenge the existing CC definition and complicates the use of a common *C. jejuni* nomenclature system based on MLST for this CC.

The spatial signal in the core genealogy, population structure, and accessory genome of ST-45 CC varied between sub-lineages, while no evidence of a temporal signal was found. The spatial signal, however, was weakly, inconclusively and variably reflected when the data was corrected for population structure, as shown in our GWAS. This supports that the observed genetic separation between Finnish and British strains is most likely due to the clonal expansion of specific clades within countries. Genomic separation between the *C. jejuni* populations in Finland and the UK has been noted before (McCarthy, Gillespie *et al.* 2012), and the authors hypothesized that the observed separation was due to differences in the epidemiology of campylobacteriosis and to seasonal fluctuations of specific CCs (i.e. ST-45 and ST-283 CCs). In our data, we found that the Finnish monophyletic branch of BAPS 2 corresponds to the expansion of monomorphic ST-230 genotype, the second most common ST in Finnish patients (Kovanen, Kivisto *et al.* 2014b) and chickens (Llarena, Huneau *et al.* 2015). In contrary, this ST is almost non-existing in British human isolates (Cody, McCarthy *et al.* 2015). This pattern of overrepresentation of certain STs according to geography was observed for the British BAPS 7 and 9, and the Baltic BAPS 10, which might reflect either an adaptation of these STs to their respective countries or the presence of an ecological barrier.

The major clade *a* consisted entirely by British strains of ST-4791, ST-7259 and ST-7256. These isolates were of non-agricultural animal origin according to the pubmlst database (pubmlst.org.net), and a major part of their core genome showed signs of recombination with lineages outside our dataset. This branch uniformly carried the three metabolic accessory genes *ggt,* fucose permease *(fucP)* and a glucose metabolism locus. The combination of *fucP* and *ggt* is rare across the species tree, while the exclusive presence of *fucP* or *ggt* is common in ST-21 CC and ST-45 CC, respectively (Gripp, Hlahla *et al.* 2011, de Haan, Llarena *et al.* 2012, Zautner, Ohk *et al.* 2012,). The relatively large genetic distance between the clade *a* and *b* and the increased r/m ratio and presence of non-typical genetic features in clade *a* indicate the lack of a shared clonal origin between these clades. This implicates that the clonal definition is unable to correctly summarize the relationship between these isolates. Moreover, a similar incongruence between the genealogy and MLST nomenclature was detected within clade *b* as well, where we frequently observed poly- or paraphyletic relationships among members of the same ST. These results complicate the implementation of a hierarchical WGS analysis using MLST in surveillance and outbreak investigations, as such an approach would deem isolates of different STs unrelated even though their genealogy shows otherwise, as seen e.g. for the ST-11 isolate (321–08). According to our analysis, this isolate was more closely related to ST-2219 than other isolates of ST-11, and such a bias could potentially lead to the wrong conclusions in a public health context.

Certain sublineages of BAPS 7 and 10 and the entire BAPS 4, 5, 6 and 11 were of mixed geographical origin, and among these, 96% of the isolates were ST-45. Genetically, BAPS 4 and BAPS 6 were surprisingly stable over time and space, and both the genetic variation (average 73 SNPs in a 1Mb non-recombining core genome) and levels of recombination were extremely limited. Moreover, our temporal investigation using BEAST and root-to-tip linear regression excluded the presence of a temporal signal within the sampling period of 12 years. Since Canadian, Italian and barnacle geese clustered within the diversity of BAPS 4 and 6, we conclude and argue for the existence of at least two globally distributed and temporally stable monomorphic genotypes of ST-45 possibly disseminated by rapid animal (migrating birds), food or human movement (French, Yu *et al.* 2014). The spread of genetically monomorphic lineages of bacteria within species of greater diversity have been documented earlier for *E. coli* O157:H7 (Leopold, Magrini *et al.* 2009) and *Salmonella enterica* serovar Typhi (Holt, Parkhill *et al.* 2008). According to Achtman (2012), the population dynamics of genetically monomorphic pathogens represent neutral evolution in the form of mild purifying selection in the place of periodic selection of fitter variants, as observed in Darwinistic evolution and for *E. coli* under laboratory conditions (Lenski, Travisano 1994). The author further suggested that the difference in evolution between the experimental and natural populations could be due to bottlenecks imposed on the genetic diversity during zoonotic and geographical transmission (Achtman 2012). The species *C. jejuni* is a zoonotic pathogen, and ST-45 is a generalist found in a myriad of agriculture animals, wild birds and mammals and is also isolated from the extra-intestinal environment such as water and sand (Sheppard, Didelot *et al.* 2013). Host jumps from animal hosts to humans as well as between animals are most probably frequent and happen at such a pace that genetic host-signature has been eroded in generalist lineages (Dearlove, Cody *et al.* 2015). Therefore, the evolutionary mechanism proposed by Achtman is certainly plausible for ST-45.

Migrating birds, food trade and people travelling could transmit and disseminate such monomorphic clones globally. However, observations of genetic variation over time would still be warranted since studies have estimated a relatively high mutation rate for *C. jejuni* with 23 to 32 SNP generated per 1Mbp per year (Achtman 2012, Wilson, Gabriel *et al*. 2009).

No temporal signal was evident in any of the sublineages, and *C. jejuni* strains isolated 12 years apart were no more different as those collected during the same year. These results indicate that monomorphic *C. jejuni* populations evolves much slower than expected (based on the prediction of Wilson et al. 2009) and suggests that the application of a single mutation rate across the species can be problematic and lead to wrong prediction of the time of the most recent common ancestor. However, further studies, both longitudinal laboratory studies and observational studies on a wide dataset of varied lineages, geographical origin and longer timespan, could possibly untangle the variability in the evolutionary speed in different *C. jejuni* lineages.

In conclusion, we identified problems with the use of a MLST based hierarchical nomenclature system for *C. jejuni* within ST-45 CC, since the WGS genealogy harbored both polyphyletic and paraphyletic STs, complicating the use of such systems in genomic epidemiology. Furthermore, we show the global occurrence and dissemination of two successful monomorphic clones of ST-45 and describe a national clonal expansion of Finnish ST-230, and predict that other monomorphic clones of *C. jejuni* will be discovered as the number of WGS studies increase. The evolutionary mechanisms and ecological conditions maintaining these clones are not known, and further research into this area is needed. The occurrence of monomorphic clones represents a problem for genomic epidemiology in surveillance and monitoring, as even WGS lacks sufficient capacity to reliably differentiate between these extremely similar isolates. Our results on occurrence of monomorphic clones among *C. jejuni* highlights the importance of two principles in genomic epidemiology; we can only exclude that isolates are epidemiologically unlinked with a considerable level of certainty and that the combination of genomic data and epidemiological data will be crucial to use WGS as a reliable and stable working tool in public health.

## Acknowledgements

The research from the INNUENDO project has received funding from European Food Safety Authority (EFSA), grant agreement GP/EFSA/AFSCO/2015/01/CT2 (New approaches in identifying and characterizing microbial and chemical hazards). J.C. and M.V. were partially funded by the COIN center of excellence. The authors wish to acknowledge CSC - IT Center for Science, Finland (www.csc.fi), for computational resources.

## Disclaimer

The conclusions, findings, and opinions expressed in this scientific paper reflect only the view of the authors and not the official position of the EFSA.

